# Contrasting biogeographic patterns of bacterial and archaeal diversity in the top- and subsoils of temperate grasslands

**DOI:** 10.1101/623264

**Authors:** Nana Liu, Huifeng Hu, Wenhong Ma, Ye Deng, Yuqing Liu, Baihui Hao, Xinying Zhang, Dimitar Dimitrov, Xiaojuan Feng, Zhiheng Wang

## Abstract

Biogeographic patterns and drivers of soil microbial diversity have been extensively studied in the past few decades. However, most research has focused on the topsoil while the subsoil is assumed to have similar microbial diversity patterns as the topsoil. Here we compare patterns and drivers of microbial diversity in the top- (0-10 cm) versus subsoils (30-50 cm) of temperate grasslands in Inner Mongolia of China along an aridity gradient covering a ~1500-km transect from arid to mesic ecosystems. Counter to the conventional assumption, we find contrasting biogeographic patterns of diversity and influencing factors for different bacterial and archaeal groups and between depths. While bacterial diversity increases with increasing aridity, archaeal diversity decreases. Microbial diversity in the topsoil is most strongly influenced by aboveground vegetation, but is most strongly influenced by historical temperature anomaly since the Last Glacial Maximum (LGM) in the subsoil. Moreover, the biogeographic patterns of top-subsoil diversity difference varies for different microbial groups and is overall most strongly influenced by soil fertility difference between depths and historical temperature anomaly. These findings suggest that diversity patterns observed in the topsoil may not be readily applied to the subsoil horizons. For the subsoil in particular, historical climate plays a vital role in the distribution of various microbial groups. Overall, our study provides novel information for understanding and predicting soil microbial diversity patterns at depth.

**IMPORTANCE:** Exploring the biogeographic patterns of soil microbial diversity is critical for understanding mechanisms underlying the response of soil processes to climate change. Using top- and subsoils from a ~1500-km temperature grassland transect, we find divergent patterns of microbial diversity and its determinants in the top-versus subsoils. Furthermore, we find important legacy effect of historical climate change on the microbial diversity of subsoil but not topsoil. Our findings challenge the conventional assumption of similar geographic patterns of soil microbial diversity along soil profiles and help to improve our understanding of how soil microbial communities may respond to future climate change in different regions with varied climate history.

## INTRODUCTION

Soil microbes play indispensable roles in the biogeochemical cycles of soil nutrients and soil formation, hence providing key ecosystem services including mediating greenhouse gases emission and climate change (1-5). Exploring the biogeographic patterns of soil microbial diversity is critical for understanding mechanisms underlying the responses of soil processes to climate change. Subsoil (i.e., soils residing > 20 cm below ground) contains more than half of soil organic carbon (OC) globally (6). Recent experimental studies have indicated that deep soils may show varied responses to global climate changes relative to the topsoil (7, 8) due to distinct soil environment, microbial assemblages and their functional responses to climate changes (9). Yet, studies on soil microbial diversity have mostly focused on the topsoil while the biogeographic patterns of microbial diversity in the subsoil across large scales remain elusive. With unique soil physical environment and microbial communities compared to the topsoil (6), subsoil may show divergent patterns of microbial diversity from the topsoil. Microbial diversity difference between top- and subsoils may also explain different soil processes and responses to global changes in top-versus subsoils (10). Hence, comparing biogeographic patterns and drivers of soil microbial diversity at different depths is important to improve our understanding of soil processes in a changing world.

Soil microbial diversity is influenced by a wide array of variables including edaphic properties (e. g. soil pH and nutrients) (11-14), vegetation (15, 16), contemporary climate (17-19) and historical climate change (20-22), etc. Compared to the topsoil, these variables may have differential controls on microbial diversity in the subsoil due to varied ranges and different orders of importance for these factors. For instance, linkages between vegetation and soil microbes can be directly mediated by plant species-specific symbioses or rhizodeposition (23). Given the predominant distribution of plant roots in surface soils, the distribution and diversity of subsoil microbes may be less affected by vegetation compared to the topsoil counterparts. Similarly, contemporary climate including precipitation (or aridity) and temperature has been shown to have a considerable effect on the topsoil microbial diversity (17-19), by restricting microbial access to soil nutrients or moisture (24, 25) and/or accelerating metabolic rates and biochemical processes (18). Such effects, however, may be dampened at depth (26) because microbial communities have a much longer turnover time in the deep soil (27, 28) and are less affected by contemporary climates.

In contrast to contemporary climate, historical climate change since the Last Glacial Maximum (LGM; the most recent glaciation, *ca*. 21 000 – 18 000 years before present) is found to be a better predictor of species richness than contemporary climate for vertebrates (29) and plants (30, 31) in Europe and North America. A recent study also suggests that climate change since the LGM may influence soil bacterial richness (20). Due to the old age of soil organic matter and long residence time of microbial communities at depth (26, 32, 33), microbial diversity in subsoil may be more strongly influenced by historical climate change compared with that in the topsoil. However, compared to edaphic and contemporary climatic factors, the effect of historical climate change on soil microbial diversity patterns remains poorly understood.

In addition to varied environmental influences, the diversity pattern along soil depth may vary among different microbial clades (34). Declining carbon substrate availability with soil depth produces oligotrophic environment at depth, which may restrict bacterial activity and promote subsurface-dwelling groups capable of utilizing recalcitrant carbon sources (35, 36). Previous studies have shown that soil bacterial diversity is typically highest in the topsoil and decreases with soil depth (34, 37). However, the diversity or abundance of different bacterial phyla may decrease (36, 37), increase (38), or remain consistent (34) along soil profile. Compared with bacteria, soil archaeal diversity patterns are much less explored at depth. Some studies reveal that the relative abundance of archaea or the archaea:bacteria ratio tends to increase with soil depth (39), while other studies assume that archaeal diversity decreases or remains constant along soil profiles (40-42). Hence, diversity variations of difference microbial groups also need to be compared to understand mechanisms driving microbial diversity patterns at different soil depths.

Here using amplicon-based sequencing of 16S ribosomal RNA genes, we compare the biogeographic patterns of bacterial and archaeal diversity in the top- (0–10 cm) versus subsoil (30–50 cm) along a temperate grassland transect in Inner Mongolia of China. As an integral part of the Eurasian steppe, this transect spans from arid to mesic ecosystems along an aridity gradient from Northeast China towards the west, covering a broad range of climate, soil physicochemical conditions and plant species richness. Coupled with a comprehensive list of edaphic, vegetation, and climatic (both contemporary climate and historical climate change) variables, we evaluate the relative importance of different environmental factors governing microbial diversity at different soil depths and the top-subsoil diversity difference. This study aims to test the following three hypotheses. (i) The biogeographic patterns of diversity vary between bacteria and archaea and among different functional groups. (ii) Microbial diversity patterns in the subsoil do not entirely mimic those in the topsoil, and the top-subsoil diversity difference varies with environmental gradients. (iii) Microbial diversity is strongly influenced by contemporary climate and vegetation in the topsoil and by historical climate change in the subsoil.

## RESULTS

### Geographic variations in soil bacterial and archaeal diversity

A survey of high-throughput amplicon sequencing for the 16S rRNA were performed to cover a large portion of bacterial and archaeal domains. According to the rarefaction results (Figure S1), curves of soil bacterial and archaeal communities almost reached an asymptote, suggesting that the sequencing depths were appropriate to survey most soil bacteria and archaea. After quality filtering, de-noising and removal of potential chimeras, a total of 3,531,946 and 4,086,723 high-quality sequences (grouping into 23,458 and 3,152 OTUs at 97% sequence similarity per sample) were obtained for bacteria and archaea, respectively.

Soil bacteria were dominated by 3 classes (*Alphaproteobacteria*, *Betaproteobacteria, Gammaproteobacteria*) and 10 phyla, including *Actinobacteria*, *Acidobacteria*, *Firmicutes*, *Bacteroidetes*, *Planctomycetes*, *Verrucomicrobia*, *Gemmatimonadetes*, *Nitrospirae*, *Chloroflexi* and *Armatimonadetes* (Figure S2). Among them, *Actinobacteria, Alphaproteobacteria*, *Acidobacteria*, *Chloroflexi*, *Nitrospirae* and *Verrucomicrobia* are predominantly oligotrophic while *Bacteroidetes*, *Gemmatimonadetes*, *Betaproteobacteria and Firmicutes* are copiotrophic (43-45). Soil archaea were dominated by three phyla, including *Crenarchaeota*, *Parvarchaeota* and *Euryarchaeota* (Figure S2). Among them, *Crenarchaeota* functions as ammonia-oxidizing archaea (AOA) (46). *Parvarchaeota* is known as acidophilic (47) while *Euryarchaeota* functions as methanogens and denitrifiers (4, 46).

The Shannon-Wiener diversity showed an overall longitudinal variation in the topsoil, decreasing for bacteria (*r* = −0.39; *p* = 0.026) and increasing for archaea from southwest to northeast (*r* = 0.58; *p* = 0.001; Figures 1 and S3). For different clades, the diversity decreased from southwest to northeast in the topsoil for the dominant oligotrophic bacterial clades (*p* < 0.05) and increased for the copiotrophic bacterial clades (*p* < 0.05; Figures 2 and S2). By comparison, the diversity of *Parvarchaeota* and rare, unclassified archaeal clades increased along the same geographic direction (*p* < 0.05) while that of *Crenarchaeota* decreased (*p* < 0.05).

**Figure 1.**
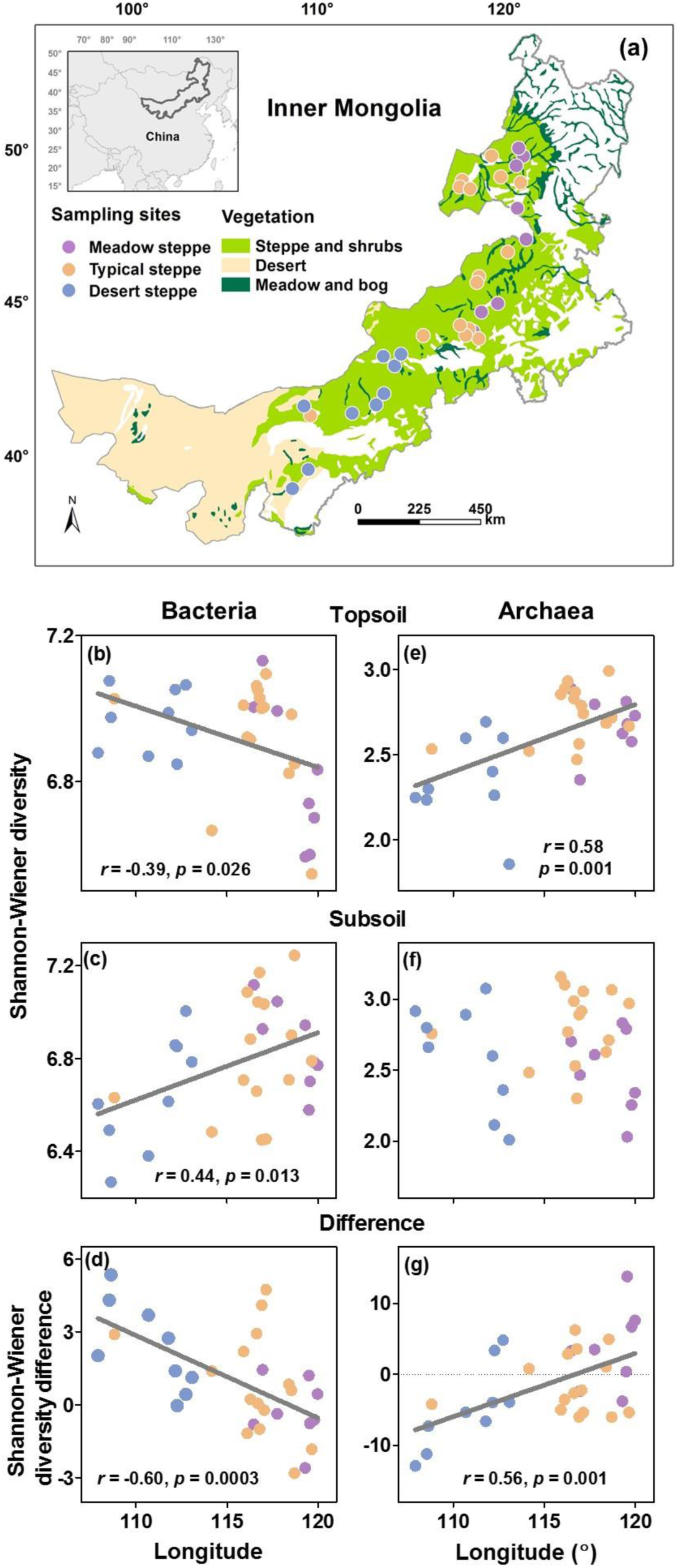
Sampling sites and geographic variation in soil bacterial and archaeal Shannon-Wiener diversity. Spatial distribution of sampling sites across temperature grasslands in Inner Mongolian grassland (a) and the changes in bacterial (b – d) and archaeal (e – g) diversity in topsoil (b, e) and subsoil (c, f) and the diversity differences (d, g) with longitude. Land cover classification is based on the Global Land Cover Characteristics Database v2.0 (https://lta.cr.usgs.gov/GLCC). Top-subsoil diversity differences in soil bacteria and archaea were estimated following Fierer et al. (10): Top-subsoil difference = (Shannon_Top_ – Shannon_Sub_) / (Shannon_Top_ + Shannon_Sub_) × 100%. Positive values of diversity differences indicate that topsoil has higher diversity than subsoil, whereas negative values indicate the opposite patterns. Solid lines indicate significant linear regressions (*p* < 0.05; n = 32).

**Figure 2.**
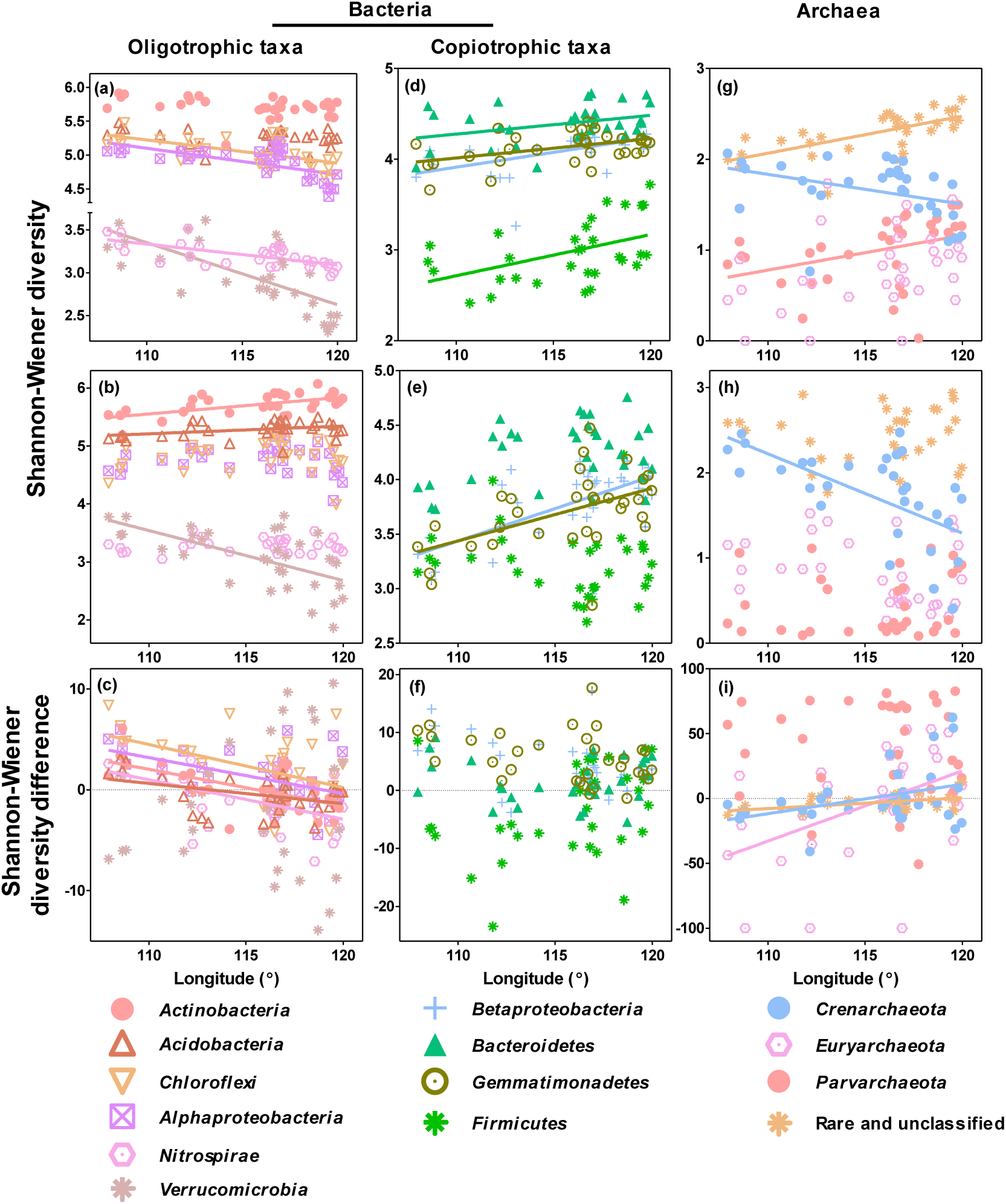
Geographic variation in the Shannon-Wiener diversity of the functional groups of soil bacteria (a - f) and archaea (g - i). (a, d, g), topsoil; (b, e, h) subsoil; (c, f, i) the top-subsoil diversity difference. For bacteria, 2 classes (*Alphaproteobateria* and *Betaproteobacteria*) and 8 dominant phyla (the rest) were categorized as oligotrophic (a, b, c) and copiotrophic (d, e, f) clades, respectively. Archaea included three phyla and some rare, unclassified clades, frequently functioning during ammonia-oxidizing (*Crenarchaeota*) and methane generation processes (*Euryarchaeota*). Solid lines indicate significant linear regressions (*p* < 0.05; n = 32).

In the subsoil, the Shannon-Wiener diversity of bacteria displayed an overall opposite geographic trend from that in the topsoil, increasing from southwest to northeast (*r* = 0.44, *p* = 0.013, Figures 1 and S3). The diversity of two oligotrophic (*Actinobacteria* and *Acidobacteria*) and two copiotrophic bacterial clades (*Betaproteobacteria* and *Gemmatimonadetes*) increased from southwest towards northeast (*p* < 0.05; Figures 2 and S2), while only one oligotrophic clade (*Verrucomicrobia*) showed a decline in species diversity in the same geographic direction (*p* < 0.05). For archaea, only the diversity of *Crenarchaeota* decreased from southwest to northeast (*p* < 0.05). The other bacterial and archaeal clades showed no consistent patterns in species diversity in the subsoil (Figures 2 and S2).

The top-subsoil diversity difference was overall positive for bacteria at the western sites and gradually decreased to negative values towards the eastern end of the transect (Figures 1 and S3). It suggests that the topsoil had a higher bacterial diversity than the subsoil in the west and had a lower bacterial diversity than subsoil in the east. This trend was mainly driven by several oligotrophic clades (*Actinobacteria, Chloroflexi*, *Alphaproteobacteria*, *Nitrospirae* and *Acidobacteria*), whereas copiotrophic bacterial clades showed no consistent trend in diversity difference (Figures 2 and S2).

In contrast, the top-subsoil diversity difference was overall negative for archaea and for some archaeal clades (*Crenarchaeota*, *Euryarchaeota* and rare, unclassified clades) at the western sites and increased to positive values towards the eastern end of the transect (Figures 1, 2 and S3), suggesting a higher archaeal diversity in the sub-than topsoil in the west and a lower archaeal diversity in the sub-than topsoil in the east. As a result, the diversity differences for bacteria and archaea were negatively correlated across the study area (*r* = −0.48, *p* = 0.0006, Figure S3).

### Explanatory variables for soil bacterial and archaeal diversity variations

The diversity of soil bacteria and archaea in topsoil, subsoil, and diversity difference displayed opposite correlations on the majority of the six groups of environmental variables, including historical temperature anomaly, contemporary climate, vegetation, soil fertility, soil pH, soil mineral (Table 1). Interestingly, the oligotrophic and copiotrophic bacterial clades in topsoil showed opposite correlations with the same variables, and so did *Crenarchaeota* and the rest archaeal phyla (Table S1). Subsoil bacterial and archaeal clades, as well as their top-subsoil diversity difference, showed no consistent correlation with environmental variables (Tables S1).

**Table 1.** Pearson correlations between soil microbial Shannon-Wiener diversity and environmental variables in the top- and subsoils and the differences between depths.

| Variables | Bacteria |  | Archaea |  |
| --- | --- | --- | --- | --- |
|  | Topsoil | Subsoil | Topsoil | Subsoil |
| **Historical temperature anomaly** | -0.28 | 0.41* | 0.53** | -0.07 |
| **Contemporary climate** | -0.52** | 0.26 | 0.57*** | 0.02 |
| **Vegetation** | -0.56*** | 0.28 | 0.61*** | 0.06 |
| **Soil fertility** | -0.47** | 0.02 | 0.19 | 0.05 |
| **Soil pH** | 0.41* | 0.10 | -0.56*** | -0.02 |
| **Soil mineral** | -0.47** | 0.24 | 0.08 | 0.06 |
| Variables | Difference |  | Difference |  |
| **Historical temperature anomaly** | -0.52** |  | 0.53** |  |
| **Contemporary climate** | -0.52** |  | 0.46** |  |
| **Vegetation** | -0.56*** |  | 0.46** |  |
| **Soil fertility difference** | -0.62*** |  | 0.51** |  |
| **Soil pH difference** | 0.24 |  | -0.24 |  |
| **Soil mineral difference** | -0.43* |  | 0.25 |  |

The one, two and three asterisks mean significant correlation at a level of $p < 0.05$ , $p < 0.01$ , and $p < 0.001$ , respectively.

Using hierarchical partitioning, we found that the environmental variables explained 45.2% and 53.2% of the variations in the topsoil bacterial and archaeal diversity, respectively (Figure 3). Among them, vegetation had the largest independent effect, explaining 11.3% and 14.8% of the variations in bacterial and archaeal diversity, respectively (*p* < 0.05; Figure 3) and showing a negative and positive relationship with bacterial and archaeal diversity, respectively (*p* < 0.05, Figure 4). Soil mineral and soil pH also had secondary and negative independent effects on the topsoil bacterial and archaeal diversity, respectively (*p* < 0.05), explaining 11.1% and 10.9% of their variations (Table 1 and Figure 3). Besides, contemporary climate showed a negative and positive independent effect on the topsoil bacterial and archaeal diversity, respectively (*p* < 0.05), explaining 8.9% and 10.6% of their variations. The other environmental variables had no significant independent effect on either bacterial or archaeal diversity.

**Figure 3.**
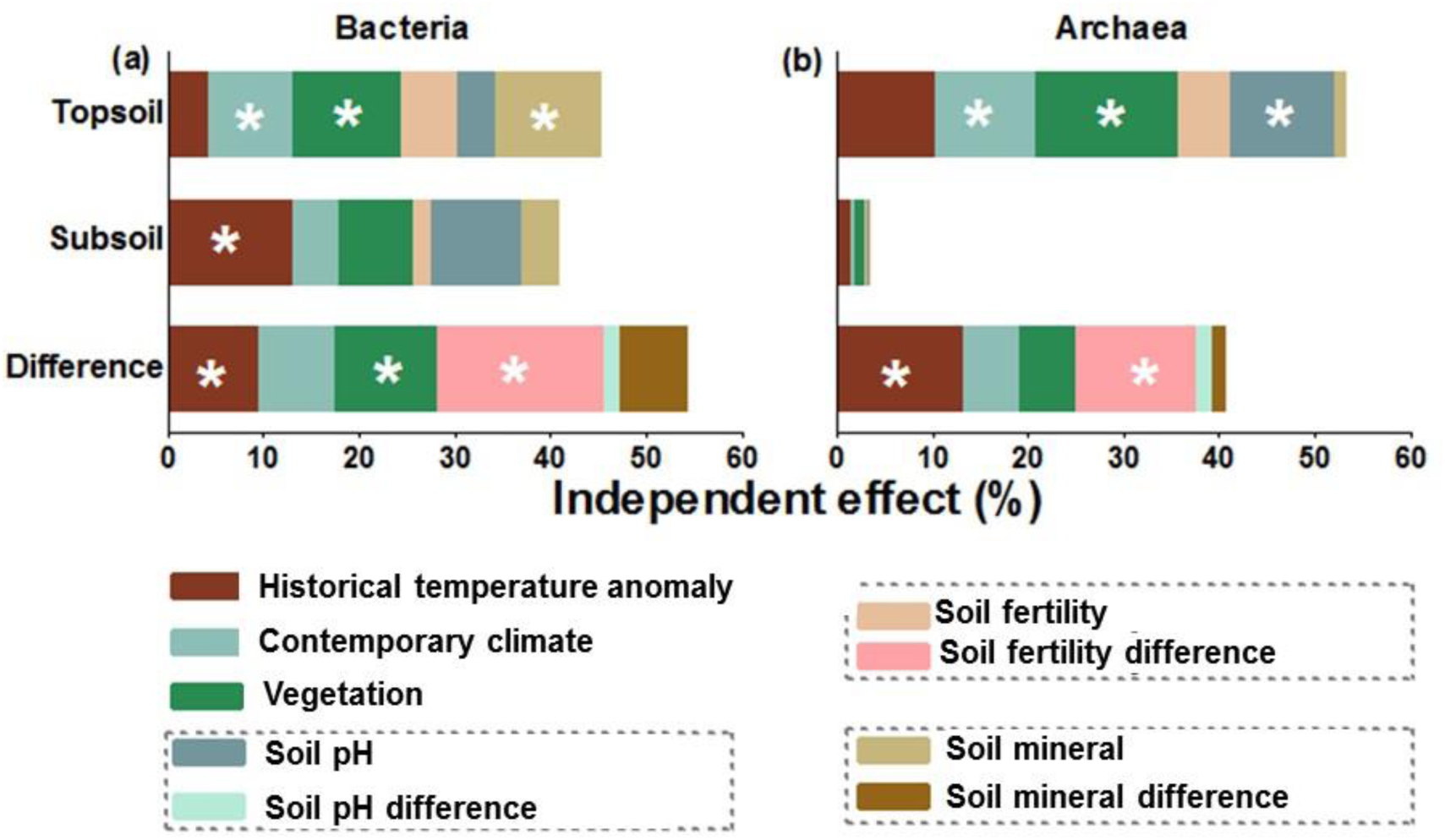
Relative importance of different environment variables on soil bacterial and archaeal Shannon-Wiener diversity. The relative importance of different environment variables was evaluated by calculating their independent effects on soil bacterial (a) and archaeal (b) Shannon-Wiener diversity and top-subsoil diversity difference using hierarchical partitioning (Table S3). Different colors of columns represented their independent effects. The asterisks indicate significant independent effects (*p* < 0.05; n = 32).

**Figure 4.**
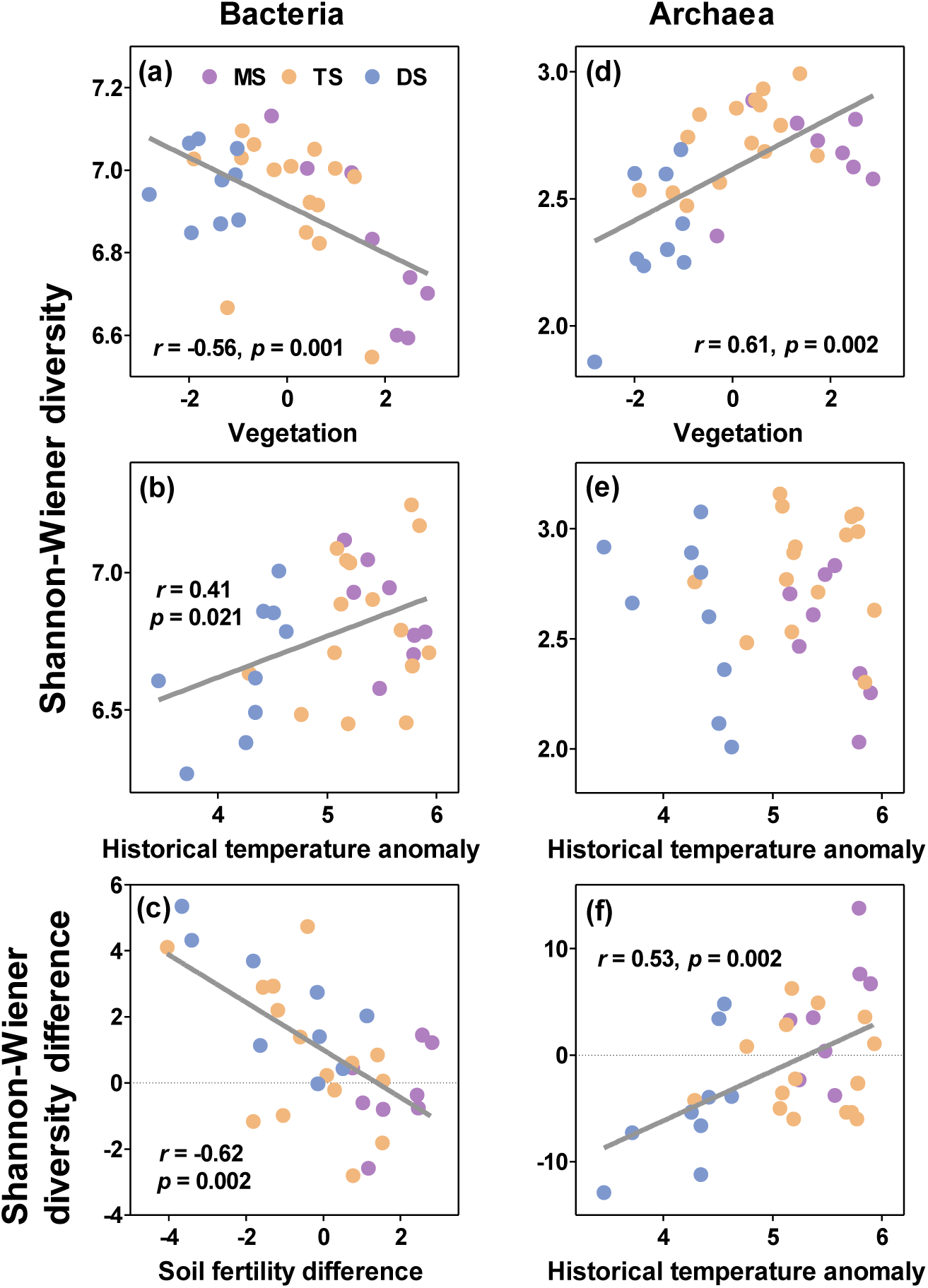
Changes in soil bacterial (a - c) and archaeal (d - f) Shannon-Wiener diversity with dominant environmental factors. (a and d), topsoil; (b and e), subsoil; (c and f), top-subsoil diversity difference. The purple, salmon and blue points indicate different vegetation types: MS (Meadow Steppe), TS (Typical Steppe) and DS (Desert Steppe). Solid lines indicate significant linear regressions (*p* < 0.05; n = 32).

In subsoil, the six environmental variables explained only 40.7% and 3.4% of the variations in bacterial and archaeal diversity, respectively (Figure 3). Among them, historical temperature anomaly had the highest positive effect on bacterial diversity (12.9%, *p* < 0.05, Figure 3), whereas other environmental variables had no significant independent effect (Figure 3). In contrast, no environmental variable exerted a significant effect on subsoil archaeal diversity, although historical temperature anomaly had the largest effect (1.3%, *p* > 0.05, Figures 3 and 4).

The biogeographic pattern in top-subsoil diversity difference was dominantly influenced by different environmental variables, distinct from that in the top and subsoils. Overall, the six environmental variables explained 54.1% and 40.6% of the variations in bacterial and archaeal diversity differences, respectively (Figure 3). Soil fertility had the highest negative independent effect on bacterial diversity difference (17.6%, *p* < 0.05), followed by vegetation (10.6%, *p* < 0.05) and historical temperature anomaly (9.3%, *p* < 0.05, Figures 3 and 4). In contrast, historical temperature anomaly had the highest positive independent effects on archaeal diversity difference (13.0%, *p* < 0.05), followed by soil fertility (12.4%, *p* < 0.05, Figure 3).

### Cascading environmental effects on bacterial and archaeal diversity

Building on the above-mentioned correlation analysis, we used SEMs to delineate causal effects of environmental variables on soil bacterial and archaeal diversity. The SEMs are developed from the priori models based on knowledge (17, 20, 48), with potential flows of causality from all categories of environmental variables to the dependent soil bacterial and archaeal diversity. The validated SEMs yield a good model fit model, indicated by a non-significant *X*^2^ test (*P* > 0.05), a high comparative fit index (CFI > 0.95), a low root mean square error of approximation (RMSEA < 0.05) (48).

In the topsoil, the constructed SEMs explained 39.7% and 47.5% of the variations in the bacterial and archaeal diversity, respectively (Figure 5a). Among the examined environmental variables, vegetation and soil mineral had significantly direct and negative role on topsoil bacterial diversity, while contemporary climate and historical temperature anomaly indirectly influenced bacterial diversity through affecting vegetation and soil mineral, respectively (Figure 5a and b). In contrast, vegetation and soil fertility had direct positive and negative effects on archaeal diversity, respectively (Figure 5a and b). Other variables only showed indirect causal effect on archaeal diversity (Figure 5a and b).

**Figure 5.**
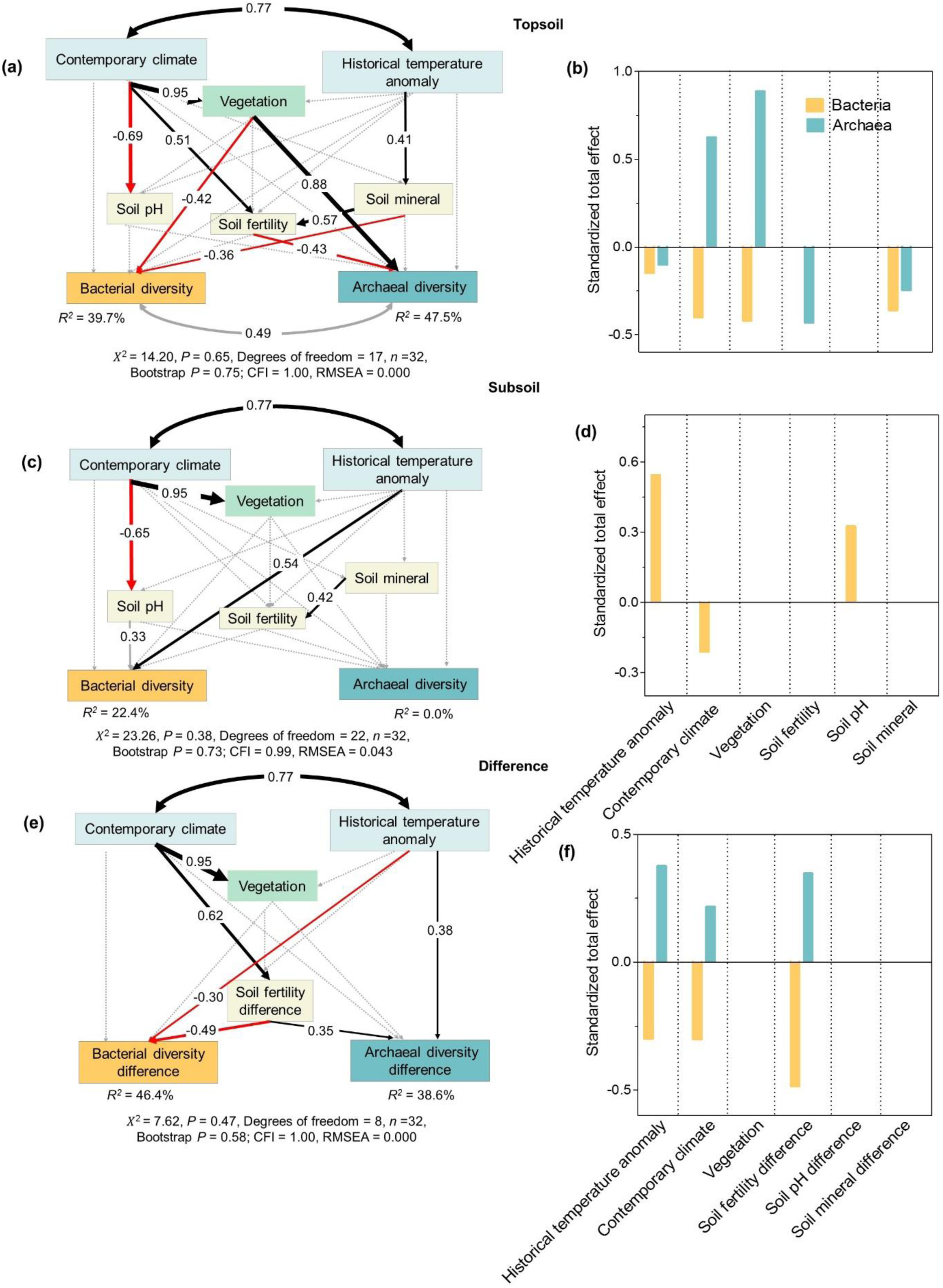
Cascading relationship between bacterial and archaeal diversity and environment variables. Structural equation model disentangling major pathways of environmental influences on soil bacterial and archaeal diversity in topsoil (a, b) and subsoil (c, d) and on top-subsoil diversity difference (e, f). Black and red arrows indicate positive and negative effects (*p* < 0.05), respectively, and their width is proportional to their standardized path coefficients (numbers on the arrows). Grey solid and dotted arrows indicate insignificant pathways from the prior models. Double-sided arrows with black and grey colors indicate Pearson correlations and modelled covariance, respectively. *R*^2^ indicates the variance of bacterial and archaeal diversity explained by the models.

In the subsoil, the constructed SEM explained 22.4% of the variations in bacterial diversity, while no significant model was yielded for archaeal diversity (Figure 5c). Historical temperature anomaly and soil pH had a direct positive effect on bacterial diversity (Figure 5c and d), although effect of soil pH is not significant (0.05 < *p* < 0.1).

The SEMs explained 46.4% and 38.6% of the variations in diversity difference of bacteria and archaea, respectively (Figure 5e). Standardized total effects of the three environmental variables, including historical temperature anomaly, contemporary climate, soil fertility difference were all opposite in bacterial and archaeal diversity difference (Figure 5f). Among six environmental variables, soil fertility difference and historical temperature anomaly had direct and negative effects on bacterial diversity difference, whereas they had direct and positive effects on archaeal diversity difference (Figure 5e and f). Other environmental variables only exerted an indirect effect on soil bacterial and archaeal diversity difference (Figure 5e and f).

## DISCUSSION

While the biogeographic pattern of soil microbial diversity has long been studied in the topsoil, its variation in the subsoil in comparison to the topsoil remains largely unknown. Here utilizing amplicon-based sequencing of 16S ribosomal RNA genes, we show contrasting microbial diversity patterns and influencing factors in the top-versus subsoils along an aridity gradient in the temperate grasslands of Inner Mongolia and reveal divergent top-subsoil diversity differences among different microbial phyla and functional groups at the regional scale.

### Divergent diversity patterns for various microbial groups in the topsoil

Our study reveals divergent geographic patterns of bacterial and archaeal diversity in the topsoil along the aridity gradient in the temperate grasslands of Inner Mongolia, with bacterial diversity decreasing and archaeal diversity increasing from southwest towards northeast (Figures 1 and S3), corroborating our first hypothesis. In all the examined environmental factors, vegetation has the strongest yet opposite effects on bacterial and archaeal diversity. Furthermore, contemporary climate including MAP, MAT and aridity index has an indirect and minor influence via its effects on vegetation (26). This result is somewhat in line with the previous findings that vegetation and contemporary climate exert a strong control on topsoil bacterial Shannon diversity shifts in arid and semi-arid temperate grassland ecosystems (19), but surprisingly towards opposite direction. The underlying reason might be because bacterial communities in the topsoil of the moisture- and N-limited Inner Mongolian grasslands (43, 49) are dominated by oligotrophic clades (such as *Actinobacteria* and *Chloroflexi*; Figure 2) due to their higher substrate affinities relative to copiotrophic clades (45). Therefore, patterns in the diversity of oligotrophic clades and their response to vegetation and contemporary climate may have dominated patterns of total bacterial diversity and environmental response.

Among the variables reflecting the effects of vegetation, plant species richness has a particularly strong negative effect on bacterial diversity (Table S2). This result stands in contrast with previous studies that reported either a positive correlation between bacterial diversity and plant diversity in the Rocky Mountains in Colorado, USA (16) or a neutral relationship in global temperate grassland (15, 50). For clades with different functions, the diversity of most oligotrophic clades is negatively correlated with plant species richness, consistent with previous studies (43, 45). In contrast, the diversity of most copiotrophic clades (such as *Betaproteobacteria*) is positively correlated with plant species richness (Table S1) (16, 50). Those results further corroborate the dominance of oligotrophic bacterial clades in infertile grassland.

In contrast to bacteria, NPP and aboveground biomass rather than plant species richness among the variables reflecting the effects of vegetation dominate the geographic patterns of archaeal diversity (Table S2). This suggests that plant carbon inputs lead to an increase in soil archaeal diversity along the aridity gradient (13, 39). Various archaeal phyla contribute differently to the diversity pattern of total archaea. Specifically, the increase of archaeal diversity from southwest towards northeast is dominated by *Parvarchaeota* and some rare, unclassified clades. The diversity of *Parvarchaeota*, which can degrade multiple carbon resources (e.g., starch, cellulose, disaccharides) (47), is positively correlated with NPP and aboveground biomass (Table S1). By comparison, *Euryarchaeota* are predominantly methanogens and capable of autotrophic growth, conferring them a relative independence from plant carbon inputs indicated by NPP and aboveground biomass (4, 46). In contrast, the diversity of *Crenarchaeota* displays a negative correlation with NPP and aboveground biomass. As *Crenarchaeota* are capable of mixotrophic growth and assimilating carbon from oxidized inorganic compounds, i.e., carbon dioxide (CO_2_) or bicarbonate (HCO_3_^−^) (4), they may compete strongly with plants for N in these N-limited grasslands, thus constraining their diversification under elevated plant growth.

In addition to vegetation, edaphic factors including soil mineral, fertility, soil pH and contemporary climate also influence the diversity of bacteria and archaea, respectively. Soil mineral negatively influences bacterial diversity, mainly due to the negative response of oligotrophic bacterial clades (Table S1). Soil fertility shows causal influence on archaeal diversity in the SEM despite their non-significant correlations with archaeal diversity. Hence, its effect primarily reflects the indirect effects of contemporary climate, historical temperature anomaly and soil mineral on archaeal diversity. Among the variables within the soil fertility group, soil total nitrogen is particularly important for archaeal diversity (Table S2), consistent with a study in Eastern China forests (51). The negative effects of soil fertility and mineral on archaeal diversity are mainly attributed to the negative response of *Crenarchaeota* (Table S1), which frequently function as ammonia-oxidizing archaea (46). Soil pH negatively influences archaeal diversity, mainly due to the response of acidophilic archaea clades, e.g., *Parvarchaeota* (Table S1) (47). This finding is consistent with previous studies on the drivers of archaeal diversity in other ecosystems (40, 52). For example, in trophic and temperate biomes across central and southern Malay Peninsula, Northern Borneo, Korea and Japan, total archaeal diversity was negatively correlated with soil pH (52). In agricultural and forest systems, thaumarchaeotal diversity was most negatively affected by soil pH (40).

### Contrasting patterns and drivers of microbial diversity in the sub- versus topsoil

Our study demonstrates contrasting geographic patterns of microbial diversity in the sub- versus topsoil of the Inner Mongolian grasslands. While bacterial diversity in the topsoil decreases from southwest towards northeast, it increases in the subsoil. Similarly, archaeal diversity shows opposite patterns in the top-versus subsoil. These results support our second hypothesis and suggest that we cannot infer the biogeographic patterns of microbial diversity in the subsoil from those in the topsoil. Furthermore, as recent studies indicate that subsoil biogeochemical processes may be strongly influenced by global climate change (7-9), it is urgent to further explore patterns as well as drivers of microbial diversity in the subsoil.

Using different statistical analyses, we find that microbial diversity is influenced by different variables in the top- and subsoil. In contrast to vegetation’s dominant control on bacterial diversity in the topsoil, historical temperature anomaly is the strongest driver of bacterial diversity in the subsoil (Figure 3), supporting our third hypothesis. Influences of historical temperature anomaly on bacterial diversity could be associated with its direct legacy effect on the distribution of soil bacteria during the past and/or indirect effects on soil mineral, soil pH, organic matter and nutrient availability. Andam et al. (53) and Martiny (21) argued that the signature of climate conditions more than 10,000 year ago could be found in contemporary soil bacterial population, such as *Streptomyces*. Previous studies also suggested that historical climate (e.g. precipitation) could affect bacterial diversity directly via its influence on enzyme sensitivity (54), or indirectly via its influence on soil properties (such as carbon stocks and quality) (20, 33, 55). For example, Delgado-Baquerizo et al. (20) found that palaeoclimatic conditions in the LGM and mid-Holocene explained more variations in bacterial richness than contemporary climate and explained a unique proportion that cannot be explained by geographic location, contemporary climate, soil properties or plant diversity.

More importantly, we show historical temperature anomaly directly regulates subsoil rather than topsoil bacterial diversity. Compared with the topsoil, microbial population and organic matter have a much slower turnover and longer residence time in the subsoil (27, 28), potentially rendering them less susceptible to contemporary than historical climate variations. Historical temperature anomaly influences subsoil bacterial diversity mainly via its effects on the diversity of *Actinobacteria*, *Acidobacteria*, *Bacteroidetes*, *Gemmatimonadetes*, *Betaproteobacteria*, and *Verrucomicrobia*, indicating that those clades may have experienced higher variability during last glacial period (20, 21).

In contrast to bacteria, no environmental factors can significantly explain the geographic patterns of subsoil archaeal diversity. However, the diversity of the biggest phylum in archaea, *Crenarchaeota,* is negatively affected by historical temperature anomaly, suggesting that this phylum might be more vulnerable to past climate variations (21, 53).

### Drivers of top-subsoil microbial diversity differences

Given the contrasting microbial diversity patterns in top-versus subsoils, we analyze patterns of top-subsoil diversity difference among different microbial groups. The diversity difference shows the opposite geographic patterns for bacteria and archaea (Figure 1), corroborating our second hypothesis. For bacteria, it decreases from positive values in the southwest to negative values in the northeast, suggesting that bacterial diversity is relatively higher in the top- than subsoil of arid and semi-arid grasslands, but is relatively lower in the top- than subsoil of mesic grasslands. The archaeal diversity difference shows the opposite trend. These results do not support the previous conclusion that soil bacterial diversity is typically highest in the topsoil and decreases with soil depth (34, 37), or that soil archaeal diversity decreases or remains constant along soil depth (40-42). Instead, our results provide a complex picture of microbial diversity contrasts between top- and subsoils and suggest that there is no universal pattern in diversity changes along soil depth for various microbial groups. Instead, the vertical variation in microbial diversity may significantly vary across different regions and among different ecosystems (e.g. arid vs. mesic grasslands).

To further reveal environmental drivers for the top-subsoil diversity difference, we explored whether top-subsoil difference in soil parameters, together with other climatic and vegetation variables, contributes to the corresponding diversity difference. The SEM analysis indicates that both soil fertility difference and historical temperature anomaly significantly influence microbial diversity difference, albeit in different direction for bacteria and archaea (Figure 5). As soil fertility difference increases from arid and semi-arid grasslands in the southwest to mesic grasslands in the northeast, bacterial diversity difference decreases. Hence, soil fertility difference has the strongest and negative effect on the diversity difference of total bacteria and the dominating oligotrophic clades (such as *Actinobacteria*, *Alphaproteobacteria*). Its negative effect is due to soil fertility’s inhibiting effect on oligotrophic clades in topsoil, whereas it exerts neutral effect in subsoil due to decreased concentrations. In contrast, the diversity difference of copiotrophic clades shows no consistent response to soil fertility difference, indicating different sensitivity among bacterial clades to soil fertility changes (43, 45). By comparison, historical temperature anomaly has a relatively minor and negative effect on the top-subsoil diversity difference of bacteria, mainly through influencing subsoil bacterial diversity.

In contrast to bacteria, historical temperature anomaly is the strongest driver on the top-subsoil archaeal diversity difference. The effect of historical temperature anomaly on soil archaeal diversity difference is primarily achieved through influencing the diversity difference of *Euryarchaeota* and some rare, unclassified clades (Table S1). As members of *Euryarchaeota* are considered to be descendants of very old cell lineages, and thus are more easily influenced by historical climate change (4). Moreover, soil fertility difference plays a secondary role in driving archaeal diversity difference, indicating that archaea are less responsive to soil fertility changes compared with bacteria. The effect of soil fertility difference is dominated by its influence on the diversity difference of *Crenarchaeota* and some rare, unclassified clades (Table S1) (13, 39). Therefore, the role of some rare, unclassified clades in regulating microbial community responses should be paid more attention to in future studies (56).

## CONCLUSIONS

Our results demonstrate contrasting biogeographic patterns of diversity between bacteria and archaea in the studied temperate grasslands, highlighting varied responses of different microbial groups to environmental variations in the soil. More importantly, by comparing microbial diversity at different soil depths, we show that microbial diversity patterns in the subsoil do not mimic that in the topsoil. Until now, studies have focused primarily on microbial diversity patterns in the topsoil. Our results suggest that these studies may misrepresent the distributions and diversity variations of vast microbial communities at soil depths. It is hence essential to add a new dimension (soil depth) to our understanding of soil microbial diversity variations along spatial gradients. Furthermore, while vegetation exerts a strong impact on microbial diversity in the topsoil, historical temperature anomaly plays a more important role in regulating bacterial diversity in the subsoil. This legacy effect of historical climate change on subsoil microbial diversity needs to be considered to better understand and predict the impacts of future climate change on soil microbial diversity.

## MATERIALS AND METHODS

### Study area and soil sampling

Our study area spans over a ~1500-km transect ranging from arid to mesic grasslands in Inner Mongolia (107.929° E ~ 119.970° E, 39.154° N~49.618° N) with varied climatic, edaphic and vegetation conditions (Supplementary Data 1, Figure S4). This transect includes several vegetation types (desert steppe, typical steppe and meadow steppe) with increasing mean annual precipitation (MAP: 165.0 ~ 411.5 mm) and decreasing mean annual temperature (MAT: 6.4°C ~ −2.3°C) from southwest towards northeast. The desert steppe is arid and low in plant species richness, dominated by perennial drought-adaptive species including *Stipa klemenzii* and *S. breviflora*, etc (57). The typical steppe has the highest coverage in Inner Mongolian with intermediate levels of net primary productivity (NPP) and plant species richness, dominated by *S. grandis*, *S. krylovii*, and *Artemisia frigida*, etc (49). The meadow steppe has the highest NPP and plant species richness, dominated by *S. baicalensis* and *Leymus chinensis*, etc (49). Soil types along this transect include Calcisols, Kastanozems and Calcic Chernozem from southwest towards northeast (49).

Soil samples were collected from 32 randomly selected sites along the transect in August 2015. At each site, five subplots (1 m × 1 m) were set at the four corners and middle of a large plot (10 m × 10 m). Three subplots along the diagonal were randomly selected for each large plot. Within each subplot, three soil cores were taken by excavating soils from predetermined depths to a total of 100 cm using a 50-mm diameter soil auger (58). Soils from the same depth and subplot were thoroughly mixed as a composite sample and divided into two portions. One portion was kept in an ice box and stored at −80°C immediately after transporting to the laboratory for DNA analysis while the other portion was air-dried for physicochemical analyses. In this study, only the topsoil (0-10 cm) and subsoil (30-50 cm) samples were used and three subplot replicates were thoroughly mixed to constitute a representative sample at each site. All soils were sieved through a 2-mm mesh with visible roots removed before laboratory analysis. The aboveground biomass (AGB) of each species was harvested by clipping the entire aboveground part, dried at 75°C to a constant weight and weighed separately for each subplot. NPP of each site was estimated using data from the Numerical Terradynamic Simulation Group (NTSG) with a spatial resolution of 1 × 1 km (http://www.ntsg.umt.edu/data).

### Soil physicochemical analysis

Total carbon (TC) and total nitrogen (TN) concentrations of soil samples were measured by combustion using an elemental analyser (Vario EL III, Elementar, Hanau, Germany). Soil OC was calculated as total carbon subtract inorganic carbon, which was analyzed volumetrically by reaction with hydrochloric acid as previously described (59). Total phosphorus (TP) was extracted using perchloric acid-sulfuric acid (HClO_4_-H_2_SO_4_) digestion and measured by colorimetric method with molybdenum blue (60). Soil pH was measured using a soil:water ratio of 1:2.5 (*w*:*v*). Soil texture was examined by laser diffraction using Malvern Mastersizer 2000 (Malvern Instruments Ltd., UK) after removal of organic matter and calcium carbonates (59). Dithionite-extractable iron (Fe_d_) and aluminum (Al_d_) were extracted from soil using the citrate-bicarbonate-dithionite (CBD) method (61, 62) and subsequently determined on an inductively coupled plasma-atomic emission spectrometer (ICP-AES, ICAP6300, Thermo Scientific, USA).

### DNA extraction and high-throughput amplicon sequencing

DNA was extracted from soils using the MoBio PowerSoil DNA isolation kit (MoBio Laboratories, Carlsbad, CA, USA) according to the manufacture’s protocol. DNA concentration was first assessed on 1% agarose gels and a NanoDrop 2000/2000C (NanoDrop, Germany) based on 260/280 and 260/230 nm absorbance ratios. According to the concentration, DNA was diluted to 1ng μl^-1^ using sterile water to serve as a template solution.

For bacteria, the V4 region of 16S rRNA gene was amplified with the forward primer 515F (5’-GTGCCAGCMGCCGCGGTAA-3’) and the reverse primer 806R (5’-GGACTACHVGGGTWTCTAAT-3’) generating a *ca*. 253 bp fragments (10). The primers contain a pair of 6-bp error-correcting forward and reverse barcode sequences, respectively. For archaea, 16S rRNA gene were amplified with a primer pair 1106F (5’-TTWAGTCAGGCAACGAGC-3’) and 1378R (5’-TGTGCAAGGAGCAGGGAC-3’) with a pair of 8-bp forward and reverse barcode sequences, generating a ca. 280 bp fragments (63). The primer set of 1106F/1378R was mainly targeted methanogenic archaeal 16S rRNA genes, but can still detect non-methanogenic clades due to non-specificity (64). All the barcodes were unique to every soil sample.

The PCR reaction was motivated in 30 μL reaction systems after mixing 15μL of Phusion® High-Fidelity PCR Master Mix (New England Biolabs), 0.2 μM of forward and reverse primers labelled with specific barcodes, and about 10 ng template DNA. Thermal cycling was repeated following the procedure: initial denaturation at 98°C for 1 min, followed by 30 cycles of denaturation at 98°C for 10 s, annealing at 50°C for 30 s, and elongation at 72°C for 30 s with a final step of 72°C for 5 min. At the termination of thermal cycling, PCR products were mixed with same volume of 1 × loading buffer (contained SYB green) and used to conduct electrophoresis on 2% agarose gel for detection. Samples with bright main strip between 400-450 bp were chosen for further experiments. PCR products were mixed in equal density ratios and then mixture PCR products were purified with GeneJET Gel Extraction Kit (Thermo Scientific). Equal molar concentrations of PCR products for each sample were pooled together. Sequencing libraries were generated using Illumina TruSeq DNA PCR-Free Library Preparation Kit (Illumina, USA) following manufacturer’s recommendations and index codes were added. The library quality was assessed on the Qubit@ 2.0 Fluorometer (Thermo Scientific) and Agilent Bioanalyzer 2100 system. At last, the libraries were sequenced on an Illumina HiSeq2500 platform and paired-end reads were generated in fastq or fasta format with forward and reverse directions assigned into separated files.

### Processing of sequencing data

Raw DNA sequences generated from the Illumina HiSeq2500 platform were processed on the Galaxy pipeline in Metagenomics for Environmental Microbiology (http://mem.rcees.ac.cn:8080/root/index) (65) at the Research Center for Eco-Environmental Sciences, Chinese Academy of Sciences. Detailedly, the raw DNA sequences assigned to samples were first cleaned by removing the barcodes and primer sequences. Then the paired-end reads were merged by FLASH (version 1.0.0), a very fast and accurate analysis tool which is designed to merge paired-end reads (66). The minimum required overlap length of paired-end reads was set at least 30 bp and the maximum overlap length approximated 90% of read pairs. The maximum allowed ratio of the number of mismatches and the overlap length was set as 0.25, the phredOffset representing the quality values of bases as 33 and the standard deviation as 10% of the average fragment length. After the mergence of paired-end reads, the sequences were filtered with the Btrim program with the threshold of average quality score > 20 over 5-bp window size and minimum length as 200 bp (67). The sequences were further de-noised by removing the sequences less than 200 bp or with ambiguous bases. Finally, the sequences were trimmed to keep sequences for bacteria between 245 and 260 bp, and for archaea between 272-288 bp, respectively, followed by exclusion of putative chimeric sequences. Therefore, we obtained a total of 3,531,946 and 4,086,723 high-quality sequences, which were grouped into 23,458, and 3,152 OTUs for soil bacteria and archaea at 97% sequence similarity and corresponding fasta format sequences were obtained using the UPARSE pipeline (68).

The reads of OTUs were annotated by referring to the Greengene database (69) for taxonomic information of bacteria and archaea with minimal 50% confidence score. Because the 505F/806R primers can target a small quantity of archaea due to its marginal non-specificity, therefore we removed the OTUs that were annotated to archaea in the following analysis. As well, the OTUs annotated to bacteria by the 1106F/1378R primer set targeting archaea were also removed.

To build a phylogenetic tree with the fasta sequences, MAFFT software (70) was first used to align the sequences, and a maximum likelihood (ML) tree was build using ExaML software (71) for soil bacteria, and RAxML software (72) for soil archaea. Both ExaML and RAxML were obtained from https://cme.h-its.org/exelixis/software.html. To make it comparable among different sites, we standardized the OUTs table across all the samples to 32,885 and 50,347 sequences (all were the smallest number of sequences across the sample) for bacteria and archaea per sample, respectively. All the following analyses were based on the standardized data. By random sampling and generating the rarefaction curves, we found that the rarefaction curves of all samples for soil bacterial and archaeal OTUs were marginally level off at the current sequencing depth.

### Climate data

To evaluate the effect of contemporary climate on soil microbial diversity, we used mean annual precipitation (MAP, mm), mean annual temperature (MAT, °C) and aridity index (AI) from 1950 to 2000. These variables have been shown to be the dominant factors of both aboveground and belowground communities in the Inner Mongolian grassland in previous studies (19). The data of MAP and MAT with a spatial resolution of 30 arc seconds were obtained from the WorldClim website (http://worldclim.org/version2) (73). AI is calculated as the ratio of MAP to potential evapotranspiration (PET). The PET data with a spatial resolution of 30 arc seconds was obtained from the CGIAR-CSI Global PET database (www.cgiar-csi.org/data/global-aridity-and-pet-database) (74).

To calculate the climate data of a site with given longitude, latitude and altitude, we conducted the following steps. First, the grid cells of a data layer within 100 kilometers from the site were extracted. Second, the longitude and latitude of the centroids of these grid cells were calculated, and their altitudes were extracted from the GTOPO30 digital elevational model with a resolution of 1 × 1 km (http://eros.usgs.gov/#/Find_Data/Products_and_Data_Available/gtopo30_info) using their centroid coordinates. Third, the following model was established for each variable respectively, using the extracted climate data, longitude, latitude and altitude of these grid cells:

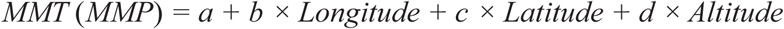

where *a, b, c* and *d* were regression coefficients. Fourth, the value of each variable at the focal plot were calculated respectively by inputting its longitude, latitude and altitude into the corresponding model. MMT, mean monthly temperature. MMP, mean monthly precipitation.

To evaluate the effect of historical climate change on soil microbial diversity, we calculated anomaly of mean annual temperature (T anomaly) as contemporary mean annual temperature minus that at the Last Glacial Maximum based on MIROC (Model for Interdisciplinary Research on Climate) (75).

### Statistical analysis

Shannon-Wiener diversity was used to estimate the diversity of OTUs (operational taxonomic unit) in topsoil and subsoil microbial communities using ‘vegan’ package in R software (version 3.4.3) (76). Differences in soil bacterial and archaeal diversity and soil properties between paired top- and subsoil were calculated following Fierer et al. (10):

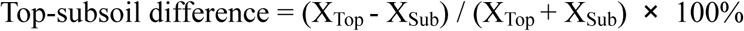

where X represents the Shannon-Wiener diversity or soil properties and the subtitles refer to top- or subsoils.

To explore the drivers of microbial diversity variations and top-subsoil diversity differences, 19 variables were compiled or measured, including historical temperature anomaly since LGM, MAP, MAT, aridity index, plant aboveground biomass, plant species richness, NPP, soil total nitrogen, soil total carbon, soil organic carbon, soil total phosphorus, soil pH, soil extractable Ca, soil extractable Mg, soil extractable Fe, soil extractable Al, soil clay, soil silt and soil sand (Supplementary Data 1, Table S2, Figures S4, details in Appendix). To avoid collinearity between variables in the following regression analysis, we classified all parameters into six groups based on their ecological implications: 1) soil fertility (including soil total nitrogen, soil total carbon, soil organic carbon and soil total phosphorus); 2) soil pH; 3) soil mineral (including soil silt, soil sand, soil extractable Fe and soil extractable Al); 4) vegetation (including plant aboveground biomass, plant species richness and NPP); 5) contemporary climate (including MAP, MAT and aridity index), 6) historical temperature anomaly. Principal component analysis (PCA) was conducted for each group encompassing more than one variable, and the first principal component (PC 1) was extracted to represent each variable group. These components explained 62.2% - 90.7% of the variations of the original variables (Table S3). PCA’s feasibility was checked using Kaise-Meyer-Olkin (KMO) test and Bartlett test of sphericity (BS) (Table S3), which indicates that PCA is appropriate to use for our data (48).

Relationships of microbial diversity and diversity difference with environmental variables were assessed by simple Pearson correlation using the R package ‘Hmisc” (77). To further compare the independent effects of different environmental factors, we conducted hierarchical partitioning using the R package ‘hier.par’ (78). The relative independent effects referred to their independent effects in the total variations.

Structure equation models were used to evaluate the direct and indirect effects of environmental factors on microbial diversity and top-subsoil differences in diversity (17). The SEMs were fitted by maximum likelihood estimation using AMOS 17 (17). For the categories of environmental variables, the PC 1 of the four variable groups and two individual variables (historical temperature anomaly and soil pH, which were standardized) were used as predictors. The prior models were evaluated and optimized by step-wise exclusion of variables with non-significant regression weights and step-wise inclusion of additional correlations based on modification indices and goodness of fit for the initial model (48). Due to our relatively small dataset with non-normal distribution, the models were modified with Satorra-Bentler correlation to improve the chi-square approximation of goodness-of-fit test statistics and confirmed using the Bollen–Stine bootstrap test (48). Models were considered to have a good fit when the bootstrap *P* value is within 0.1-1.0. Since there is no single universally accepted test of overall goodness of fit for SEMs, we also used χ^2^ test and the root mean square error of approximation (RMSEA) as criteria to text the goodness of the model fit (17). The model has a good fit when χ^2^ is low and RMSEA is near 0 (17). We checked the bivariate relationships between all variables to ensure that a linear model was appropriate (Figure S5).

## Data Availability

Hiseq2500 sequencing data will be available online upon request.

## SUPPLEMENTARY MATERIAL

Appendix.

Table S1-S3.

Figure S1-Figure S5.

Dataset: Supplementary Data 1.

## ACKNOWLEDGEMENTS

This work was funded by the National Key Research Development Program of China (2017YFA0605101, 2015CB954201), the National Natural Science Foundation of China (31621091), the State Key Laboratory of Vegetation and Environmental Change (Grant No. LVEC Y7206F2001), the Youth Fund of Ministry of Education Laboratory for Earth Surface Processes of Peking University (Grant No. LESP201702), and the Chinese Academy of Sciences-Peking University Pioneer Collaboration Team.

## AUTHOR CONTRIBUTIONS

Z.W., X.F. and N.L. designed the research. W.M., H.H., Y.L. and B.H. collected soil samples. N.L. and X.Z. conducted microbial and chemical analyses with Y.D. contributing the analytical platform for sequencing data. D. D. built the ML tree of bacteria. N.L. and Z.W. performed data analyses. N.L., X.F. and Z.W. wrote the paper with inputs from all other coauthors.

